# Linking big biomedical datasets to modular analysis with Portable Encapsulated Projects

**DOI:** 10.1101/2020.10.08.331322

**Authors:** Nathan C. Sheffield, Michał Stolarczyk, Vincent P. Reuter, André F. Rendeiro

## Abstract

Organizing and annotating biological sample data is critical in data-intensive bioinformatics. Unfortunately, metadata formats from a data provider are often incompatible with requirements of a processing tool. There is no broadly accepted standard to organize metadata across biological projects and bioinformatics tools, restricting the portability and reusability of both annotated datasets and analysis software. To address this, we present Portable Encapsulated Projects (PEP), a formal specification for biological sample metadata structure. The PEP specification accommodates typical features of data-intensive bioinformatics projects with many samples, whether from individual experiments, organisms, or single cells. In addition to standardization, the PEP specification provides descriptors and modifiers for different organizational layers of a project, which improve portability among computing environments and facilitate use of different processing tools. PEP includes a schema validator framework, allowing formal definition of required metadata attributes for any type of biomedical data analysis. We have implemented packages for reading PEPs in both Python and R to provide a language-agnostic interface for organizing project metadata. PEP therefore presents an important step toward unifying data annotation and processing tools in data-intensive biological research projects.

## Introduction

Biological data generation is accelerating, and considerable effort is now being invested in how to best share it. These efforts include expansions of databases^1,2^ as well as new data standards and ontologies, including the FAIR guiding principles and other guidelines for data sharing^3–8^. Major effort is being invested in building an open data ecosystem upon which data of many types may be easily shared and reused.

As our ability to measure and store data has increased across scientific disciplines, analysis has frequently become the bottleneck of scientific advance. To mitigate this, new computational pipelines and analysis approaches are under constant development. These pipelines are increasingly federated though pipeline frameworks, leading to now dozens of such frameworks that simplify developing reusable computational pipelines^9^, as well as standards for workflows such as the common workflow language^10^, SnakeMake^11^, Galaxy^12^, and Nextflow^13^. Similarly, new containerization technology is making computing environments more portable^14–16^ and efforts to build data commons^17^ and cloud analysis platforms^18^ are bringing analysis to data hosted in the cloud. Collectively, these efforts seek to meet the challenge of reproducible analysis in a complicated and growing ecosystem that combines public and private data.

Efforts to both curate open biological data and to standardize bioinformatics analysis are certainly complementary, but progress in each area independently does not necessarily make it easier to connect the two. In fact, relatively less effort has been placed at the confluence of data and analysis in biology. We may call this connection a “data interface,” which describes how a dataset connects to an analysis tool (Fig. 1A). As it stands, published bioinformatics pipelines, even if reproducibly built in a standard framework, typically describe a unique data interface, requiring a user to manually structure data repeatedly to fit each pipeline (Fig. 1B). On the flipside, data repositories also typically expose an individual procedure such as an API for accessing the data. In practice, it requires substantial manual effort to plug an arbitrary dataset into an arbitrary analysis tool – even if both adhere to best-practice community sharing and analysis development standards.

**Fig. 1:**
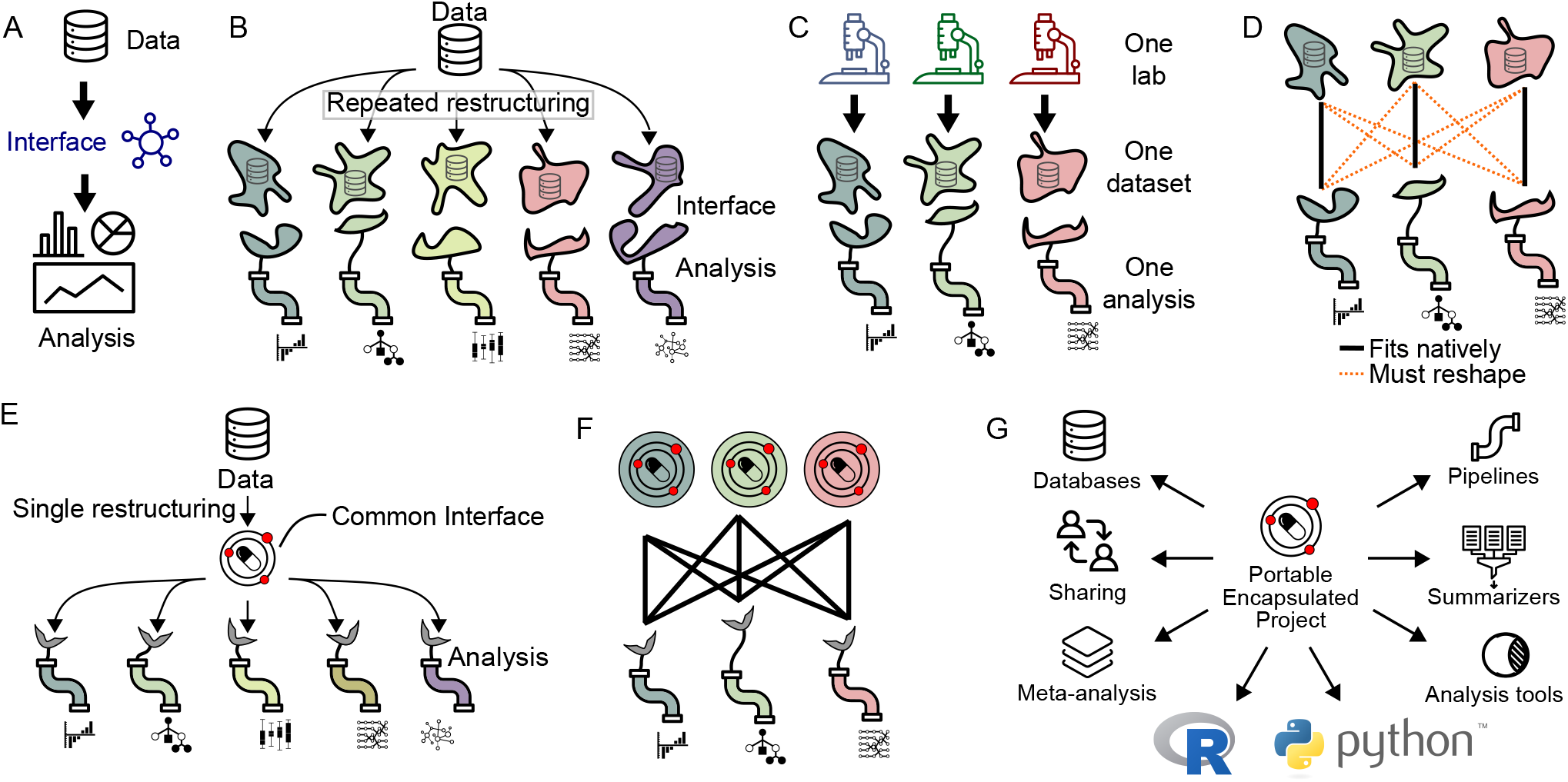
A data interface links data to analysis. A) Schematic of a data interface. B) Each analysis typically describes its own unique data interface. The one lab, one dataset, one analysis mode of research tightly couples datasets and analysis. D) With individual data interfaces, running a data set through multiple analyses requires reshaping the data for every pairwise connection of data and analysis. E) The PEP specification provides a standardized interface that reduces reshaping. F) Using PEP, no reshaping is required to run a data set through a different analytical tool. G) A PEP may be used in different contexts, and by a variety of tools and programming languages.

This challenge is surmountable for a typical project that links one data set to one analysis process – the *one lab, one dataset, one analysis* approach, which has been the dominant model (Fig. 1C). But imagine an attempt to link multiple datasets from multiple sources to multiple analysis tools. Each pair of data and tool requires a unique data description, which probably requires substantial manual data munging (Fig. 1D). The result is that analysis done by an individual lab is often restricted to a particular dataset generated by that lab for that project. What would it take to build a computing ecosystem that would relax this coupling, making it routine to mix-and-match data and pipelines across groups?

A first step to realize this vision is to standardize the data interface. This would make both datasets and tools more portable, facilitating data integration and tool comparison. To this end, we present the Portable Encapsulated Projects (PEP) specification. The PEP specification standardizes the description of sample-intensive biological research projects, enabling data providers and data users to communicate through a common interface (Fig. 1E). This standardization facilitates using different pipelines for the same datasets (Fig. 1F). In addition to standardization, the PEP specification provides powerful portability called *project modifiers* and *sample modifiers* that make project metadata annotation independent of a particular computing platform. PEP also provides a customizable validation framework that can be used to first define and then to validate the sample properties required for a particular application. Finally, we provide tools that read PEPs and handle PEP modifiers in R and Python, which can be extended by specialized tools.

PEP thus provides a unifying data organization that can be employed by many tools to make it easier to share data and tools. The goal of PEP follows the vision of the Investigation/Study/Assay (ISA) biological metadata management framework^19^. Relative to ISA, PEP emphasizes generality, programmatic metadata preprocessing, and integration into workflow systems. Existing tools can easily accommodate the PEP structure; for example, SnakeMake includes a special directive to directly import a PEP into a workflow that functions alongside earlier, specialized data formats. Similarly, our companion tool, *looper*, can be used to submit arbitrary CWL workflows to a CWL runner for each sample in a PEP project. This sets the stage for a single data description that can be used as input for multiple workflows – even workflows built using different frameworks.

Together, these advantages realize a unified specification that can be read and processed by many types of downstream analysis (Fig. 1G). By standardizing the description of project metadata and providing standalone, modular tools to read that standard, we simplify existing processes and enable new types of analysis. For instance, standardized PEPs enables meta-analysis that assesses sample properties across hundreds of projects, since each project can be read the same way. Databases, instead of requiring a custom project description and file naming and organization could instead simply provide a schema and use PEP to load published projects into a structured database. Tools that summarize processed data can be made to use the same PEP that runs the original workflows, making these kind of summarizing tools more broadly applicable. And finally, a shared metadata structure simplifies sharing across individuals and tools.

## Results

### Basic PEP specification

The *PEP specification* defines a way to organize project and sample metadata in files using `YAML` and `CSV` formats. The term *project* refers to a collection of meta-data that describes a set of samples. A *sample* is defined loosely as any unit that can be collected into a project; it consists of sample attributes, usually with one or more that point to data files. A *PEP* is a set of files that conform to the PEP specification. An common example could be a typical biological research *project* made up of a set of RNA-seq *samples* grouped to answer a particular question.

The specification defines a PEP in two files: A `YAML` configuration file, and a tabular comma-separated value (`CSV`) annotation file (Fig. 2 A). The configuration file provides project-level descriptions, such as paths to remote or local sources of data, global analysis parameters, or other project attributes. The tabular file is a sample table, providing metadata attributes for each biological specimen included in the project. An optional third file, the subsample table, can be used to specify sample attributes with multiple values (see http://pep.databio.org for further details). A basic PEP configuration file has just a few fields in `YAML` format, such as this example `YAML` file (Fig. 2B) that points to a `samples.csv` file (Fig. 2C), which contains a header line of sample attributes and then one data row per sample. Together, these two files describe a minimal project. The basic PEP format is thus extremely flexible and can accommodate assorted sample-intensive biological research project data. Because PEP uses simple plain text files, it is universally accessible, easy to version control, and inexpensive to store.

**Fig. 2:**
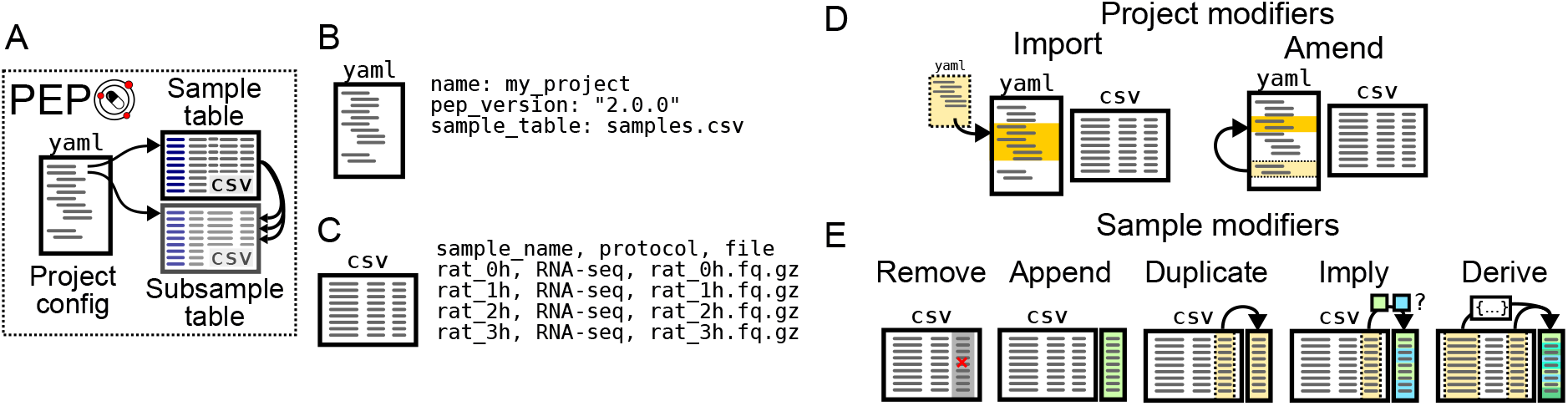
The PEP specification. A) A PEP consists of a YAML configuration file, a sample table, and a subsample table. B) The YAML file describes project-level attributes. C) The sample table (and subsample table) describe sample-level attributes. D) Project modifiers allow the PEP to import values from other PEPs, or embed multiple variations within a single PEP. E) Sample modifiers can change sample attributes by using the project config YAML file, without actually changing the CSV file.

This very simple approach is then extended in two critical improvements: First, we added features that improve portability called *project modifiers* and *sample modifiers*, which enable us to remove environment-specific file paths and analysis-specific metadata from the sample table, making it easier to use a single meta-data representation for multiple analyses in different computing environments. These *modifiers* are handled by implementations of the PEP specification, which then provide *modified*, or *processed*, sample and project metadata for downstream tools to Consume. Second, we built a validation framework for PEPs that includes a base schema to validate generic PEPs along with tools to extend this schema to more specific use cases. This generic + specialization approach allows us to construct a re-usable project definitions that can be extended modularly to provide increased specificity. Together, these two improvements provide the power and specificity that enables PEP to unify and enhance our metadata descriptions for many types of data-intensive biological research projects. We describe these in more detail below.

### Project modifiers

Project modifiers are special project attributes that provide additional functionality to a project. The two modifiers are *import* and *amend*, which allow users to either merge or embed PEPs (Fig. 2D). At times it is useful to create two projects that are very similar, but differ just in one or two attributes. For example, you may define a project with one set of samples, and then want an identical project that uses a different sample table. Or, you may define a project to run on a particular reference genome, and want to define a second project that is identical, but uses a different reference genome. You could simply define 2 complete PEPs, but this would duplicate information and make it harder to maintain. Instead, project modifiers make it easier to tie projects together through the *import* and *amend* relationships.

#### Project modifier: import

The *import* project modifier allows the configuration file to import other PEPs. The values in the imported files will be overridden by the corresponding entries in the current configuration file. Imports are recursive, so an imported file that imports another file is allowed; the imports are resolved in cascading order with the most distant imports happening first, so the closest configuration options override the more distant ones. Imports provide a way to decouple project settings so that more specific projects can inherit attributes from more general projects. Imports allow users to combine multiple files into one PEP description. The import modifier handles sample tables the same way it does any other attribute. If a sample table is specified in both an imported and importing PEP, it does not merge or update individual samples or tables, but simply selects the highest priority value of the `sample_table` attribute.

#### Project modifier: amend

The *amend* project modifier allows the configuration file to embed multiple independent projects within a single PEP. When a PEP is parsed, you may specify one or more included amendments, which will amend the values in the processed PEP. Amendments are useful to define multiple similar projects within a single project configuration file. Under the *amend* key, you specify names of amendments, and then underneath these you specify any project variables that you want to override for that particular amendment. It is also possible to activate more than one amendment in priority order, which allows you to combine different project features on-the-fly.

Example:

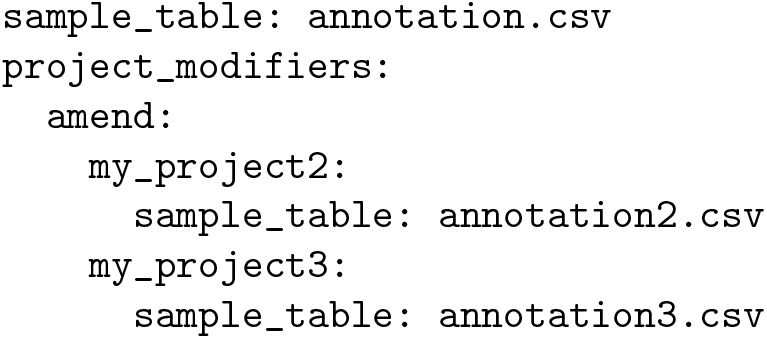

When used in tandem, imports and amendments together make it possible to create powerful links between projects and analysis settings that can simplify running multiple analyses across multiple projects.

### Sample modifiers

Sample modifiers are project-level settings that adjust sample attributes. After the sample table is read, sample modifiers are applied, adding new attributes or changing attributes from the original sample table. Sample modifiers enable keeping analysis-specific sample attributes in the project configuration file so the sample table can be more easily shared across projects. This allows the creation of a sample table that does not need to be edited when moved to either a different project or compute environment, making both project and sample metadata more portable.

You can add sample modifiers to a PEP by adding a `sample_modifiers` section to a project configuration file. Within this section, there are 5 subsections corresponding to 5 types of sample modifier (Fig. 2E). Three modifiers – *remove*, *append*, and *duplicate* – are very simple operations. The more expressive sample modifiers – *imply* and *derive* – lend considerable flexibility to the construction of PEP sample tables.

#### Sample modifier: remove

The *remove* modifier simply removes a specified attribute from all samples. It can be useful if a particular analysis needs to eliminate a particular attribute without modifying the original sample table.

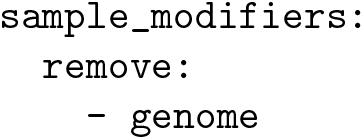

#### Sample modifier: append

The *append* modifier adds constant attributes to all samples in a project. For example, if you write `genome: hg38` as an entry under `append`, then when the PEP is parsed, the samples will each have an additional attribute, `genome`, with value `hg38`. This modifier is useful because it allows keeping static attributes in the project configuration file. It also allows you to preserve project-level information (like `genome`) separate from sample-level information, but still pass that information along to pipelines that require it for each sample. This addresses the structural mismatch in independence that follows from project composition – very often, samples may be processed independently while having high dependence among their metadata. PEPs are friendly to the *don’t repeat yourself* principle that improves project maintainability.

Example:

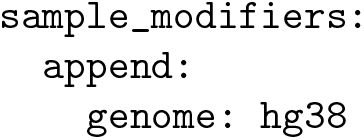

#### Sample modifier: duplicate

The *duplicate* modifier allows copying an existing sample attribute into a new one. For example, the “genome” attribute could be a synonym of the “Genome” attribute. This allows us to tweak settings at the project level, which simplifies use of an alternate pipeline with different requirements, without requiring modification of the underlying sample table that may break earlier analysis. In the `key:value` pair, the old attribute name listed as `key` will be duplicated to create a new attribute named with the corresponding `value`.

Example:

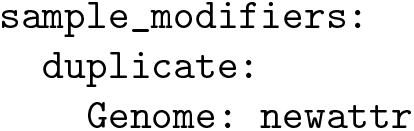

#### Sample modifier: imply

The *imply* modifier lets a user add sample attributes that are modulated based on the value of an existing sample attribute. For example, a common use case is to use *imply* to set a *genome* attribute for any sample with a specific value in its *organism* attribute. This enables complete separation of description of sample-intrinsic properties (like organism) from project-level values (like reference genome, which may change).

Example:

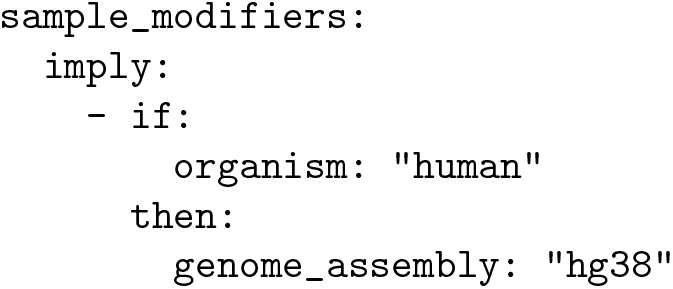

#### Sample modifier: derive

The most expressive sample modifier is called *derive*. This modifier allows us to create sample attributes that are derived from other sample attributes. The most common use case is to specify paths to data files at the project level instead of at the sample level. This allows tabular sample descriptions to avoid including any environment-specific information (such as a file path), so that moving a project from one compute environment to another requires editing only a single line in the project configuration file.

The *derive* modifier consists of two pieces of data: First, the *attributes* section lists sample attributes to be derived. Second, the *sources* section contains key-value pairs, where the keys are source names and values are string templates. The source names are the original values of the derived attributes. The string templates are used to derive new attribute values by the PEP processor, replacing the source names in the original table. These templates may contain sample attributes enclosed by curly braces, such as {`sample_name`}.

Example:

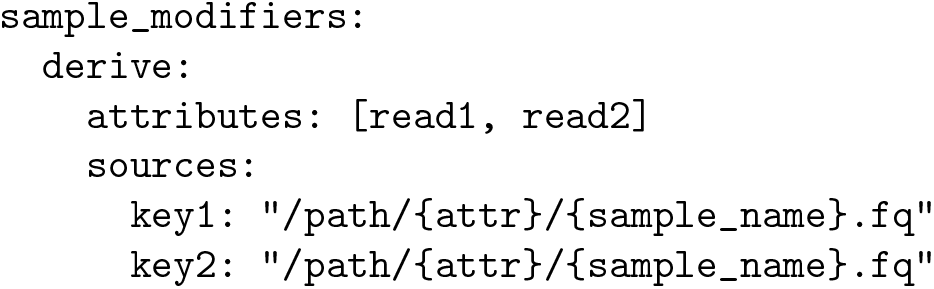

In this example, {`attr`} and {`sample_name`} represent other attributes that are present on the sample. These may be populated from the sample table, or from other attributes that have been added using a sample modifier such as `append`.

When derived source paths include a shell variable, derived attributes enable not only a sample table, but an entire PEP, to be made completely portable with no editing. For instance, we could replace /`path`/ above with `$DATAPATH`, and this PEP would then point to the correct files on any computing environment with the `$DATAPATH` environment variable set.

### Project and sample validation

To make it easier to build valid PEPs, we also implemented a PEP validation tool called `eido`. Eido is a specialized PEP validator based on JSON-schema (https://json-schema.org/). Complete documentation, descriptions of schema features, and example schemas can be found at eido.databio.org. Schema files may be equivalently saved in either JSON or YAML format. Eido can be used with a generic PEP specification schema to validate a PEP in general. Even more important, tool authors can provide a schema that describes more specific requirements for a tool, and eido can validate a given PEP to make sure it conforms to both the generic schema and the more stringent schema, ensuring that it can run on a particular tool (Fig. 3A).

**Fig. 3:**
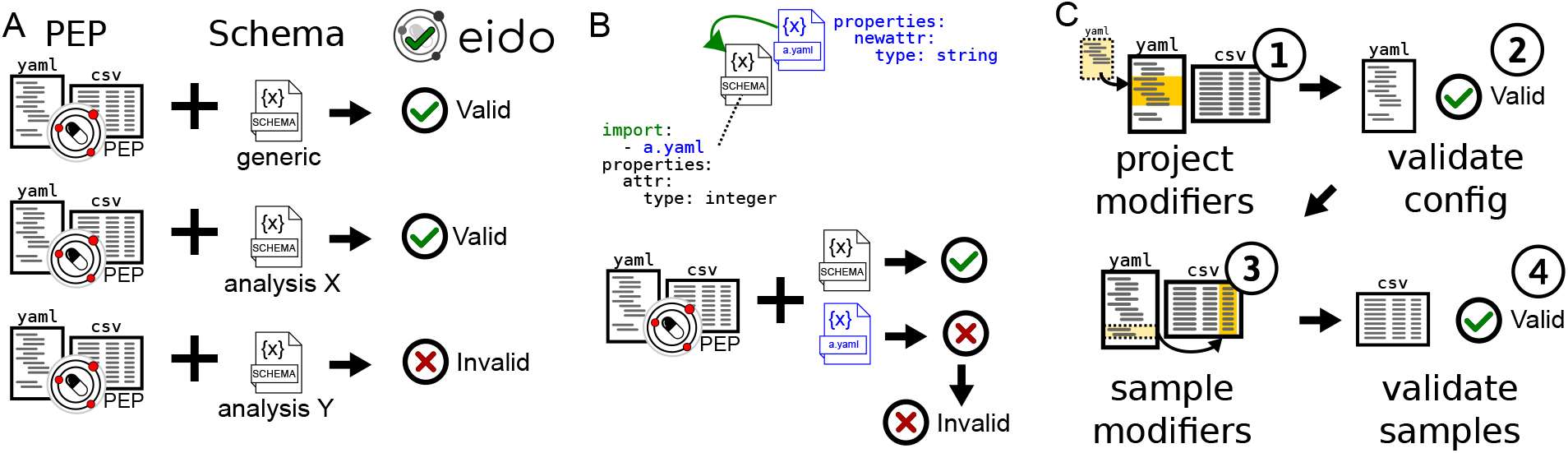
PEPs can be validated against generic or specific schemas. A) A generic schema ensures compliance with the PEP specification, while specialized schemas describe requirements for a particular analysis. B) PEP schemas can import other schemas. C) Validation uses two steps so samples are validated after PEP modification.

For example, an author of a pipeline may write a schema specifying that samples must have attributes named `read1` and `read2`, which must be of type `string`, and which must point to input files. Furthermore, the schema specifies that samples must have an attribute called `genome` that specifies the genome to align to, perhaps with a list of allowable values. With this schema published, it is now possible to validate a PEP to ensure that it fulfills the requirements for this pipeline. PEP schemas can also import other schemas (Fig. 3B). In this case, the PEP must validate against all requirements specified by imported schemas to be valid.

Specific schemas for PEPs are written using JSON-schema with a few additions that extend the basic vocabulary to tailor it to the PEP use case. For example, our validator adds the term `required_files`, which allows a schema author to indicate which sample attributes must point to files that exist. Because eido is based on JSON-schema, it inherits the explicit variable typing (*e.g.* `string`, `number`, `boolean`), and restrictions on values (*e.g.* ranges, regular expressions, enumerated values). Eido uses a two-stage validation that first validates the configuration file, and then validates individual samples *after* they have been processed (Fig. 3C). This ensures that sample attributes that are added or modified can be properly checked. These adjustments to the basic JSON-schema validation allow eido to satisfy the requirements of validating bioinformatics research projects.

### PEP implementations in R and Python

The reference implementation of the PEP specification is the `peppy` python package, available from the Python Package Index (PyPI). Peppy instantiates in-memory project objects and provides a Python API for programmatic access to any project metadata from within Python. A user simply creates a Project object (`prj` = `Project`(“`config.yaml`”)) and may now interact with the project metadata within Python. This package is a generic, extensible object framework that enables developers to build additional tools using these objects. For instance, SnakeMake relies on the `peppy` package to handle parsing and reading PEP-formatted project metadata to power a workflow run.

We have also developed an R implementation of PEP in the `pepr` package, available on CRAN. PEP files can be parsed in R with a similar function call, `prj` = `pepr::Project`(“`config.yaml`”), which provides an R API for interacting with PEPs in R. These tools provide a PEP project interface to programmers of two of the most popular data science programming languages, increasing portability of PEP projects. We and others have successfully used this infrastructure in dozens of projects with hundreds to thousands of individual samples.

We are interested in future efforts to expand this to other computing frameworks. These APIs provide basic functions for interacting with projects and samples, including setting and accessing variables, extracting the sample attributes and sub-attributes as a tabular object (using pandas in Python and data.table in R), accessing individual samples as objects. In each case, all the sample and project modifiers are processed behind the scenes so downstream tools can easily make use of the PEP portability features. The formal API is documented in the respective package documentation.

## Discussion

### The promise of PEP

As the amount of available data increases, it is useful to build a common infrastructure to link it to analytical tools. Currently, downloading and analyzing an external dataset requires significant manual investment. Because each analytical pipeline typically has a unique interface to input data, testing multiple competing pipelines on a single dataset requires describing the dataset multiple times. These manual steps hinder re-analysis and re-use of existing data.

We here propose reducing this barrier with the concept of Portable Encapsulated Projects. The PEP specification is at once standardized and flexible. It provides a loose generic specification that can be easily extended for specific use cases. It also provides a validation framework that can easily accommodate both generic and specialized PEPs.

Together, PEP provides an interface between data and tools that makes each more useful. If a tool developer designs a tool to read PEPs, then it is immediately possible to apply the tool to any published, compliant PEPs. To describe how to use the tool, the developer needs only define a PEP schema, which can be validated using `eido`; any project defining these attributes would then work without modification. Users then immediately know how to format a project for the tool, and by describing newly generated data in PEP format, they may immediately plug that project into the tool. As developers build pipelines that understand PEP format, they make it simple to apply their pipeline to new PEP-compatible projects as they emerge.

On the flipside, as data producers publish datasets in PEP format, they make it easy for pipeline developers to test new analytical techniques on data from a variety of sources. This will incentivize data sharing and re-use, driving innovation and discovery both in tool development and in understanding of data.

Together, these tools create a programmable link between data and analysis, making it simple to re-analyze an existing dataset with a newly developed pipeline, grab a relevant public dataset to include with newly generated data in a private project, or test a published PEP-compatible pipeline on some in-house data.

### PEP in practice

Several projects have been published that make use of PEP. From our research group, we have produced PEP-compatible pipelines for several biological data types, including nascent RNA sequencing^20^ and ATAC-seq^21^. We also rely on PEP for listing reference genome assets for *refgenie server*^22,23^, and we used PEP in a study of generating simulated genomic intervals^24^. These examples and others provide a starting point for interested developers or users who would like to see PEP in action. They also demonstrate the breadth and versatility of PEP.

### A call for community involvement

To conclude, we offer a call for community involvement to support reaching the vision of metadata interoperability. Three key steps will be required before this can happen: First, we need tools that support and extend the PEP specification; second, we need adoption by workflow engines; and finally, we need support of public datasets and data repositories to accept and provide data that fits the system.

A first step will be to build tools that operate in this area. To facilitate community uptake, we are developing a series of tools that subscribe to the PEP standard. Above, we described Python and R packages that read PEPs, along with `eido` for PEP validation. These core tools can form the foundation of new tools, and we hope that others in the community will use them to add functionality to the PEP ecosystem. For our needs, we are extending these capabilities with several ongoing projects: First, `geofetch` is a data fetcher that accepts a list of SRA or GEO accession numbers and then downloads raw sequence data from the Sequence Read Archive and constructs a PEP, ready to be plugged into a PEP-compatible analysis tool. Second, `looper` is a workflow-engine-agnostic command submission engine that reads PEP-formatted sample data and runs arbitrary commands. Finally, `BiocProject` is an upcoming project that adds bioconductor-specific functionality to PEPs, simplifying biological data analysis of PEPs in R.

A second step will be for workflow engines to adopt PEP as a way to specify samples. Workflow engines are becoming a critical component of biological data analysis, and as such, they provide an important incentive for the way users and tool developers organize metadata. Unfortunately, most workflow engines still require a custom format for describing input metadata. We have been reaching out to workflow engine communities, such as the SnakeMake^10^ and CWL^11^ communities, which already have some support for PEP-formatted metadata. We are also working on a conversion function in `eido` that would allow users to write custom formatters, making it easier to fit PEP-formatted metadata into custom analyses. We invite collaboration and involvement from other workflow-oriented communities who could support a community effort for standardized metadata organization that spans workflow engines.

And third, another important step will be for datasets and data repositories that understand this format, both for submission and download. We encourage authors of individual papers to consider using a PEP-structured sample table when publishing descriptions for individual projects. And we invite large-scale data providers to make it possible to download data descriptions in PEP-compatible files, and even to submit data in PEP-valid format.

To our knowledge, this is the first major effort to produce a universal specification and framework for collections of biological sample metadata geared toward meta-data and data processing. PEP can be tailored with ease to specific use cases with schemas that define specific tool requirements. We anticipate that these tools will encourage both bioinformatics pipeline developers and data producers to subscribe to a common format, benefiting both and leading to increased ability to extract useful information from biological data.

## Availability

All described software is BSD2-licensed and developed on GitHub at github.com/pepkit. The Python implementation is on PyPI and the R implementation is on CRAN. The formal PEP specification can be found at pep.databio.org.

Identifiers:

- `eido`: RRID:SCR 021076; biotools:eido-python-package
- `pepr`: RRID:SCR 021077; biotools:pepr-R-package
- `peppy`: RRID:SCR 021078; biotools:peppy-python-package

## Acknowledgments

We thank Johannes Köster, Jason Smith, Aaron Gu, and the Sheffield lab for input. This work is funded by the National Institutes of Health Institute for General Medical Sciences (NIGMS) award R35GM128636 to NCS.

